# A simple explanation for declining temperature sensitivity with warming

**DOI:** 10.1101/2021.01.12.426288

**Authors:** E. M. Wolkovich, J. L. Auerbach, C. J. Chamberlain, D. M. Buonaiuto, A. K. Ettinger, I. Morales-Castilla, A. Gelman

## Abstract

Temperature sensitivity—the magnitude of a biological response per °C—is a fundamental concept across scientific disciplines, especially biology, where temperature determines the rate of many plant, animal and ecosystem processes. Recently, a growing body of literature in global change biology has found temperature sensitivities decline as temperatures rise (Fu et al., 2015; Güsewell et al., 2017; Piao et al., 2017; Chen et al., 2019; Dai et al., 2019). Such observations have been used to suggest climate change is reshaping biological processes, with major implications for forecasts of future change. Here we present a simple alternative explanation for observed declining sensitivities: the use of linear models to estimate non-linear temperature responses. We show how linear estimates of sensitivities will appear to decline with warming for events that occur after a cumulative thermal threshold is met—a common model for many biological events. Corrections for the non-linearity of temperature response in simulated data and long-term phenological data from Europe remove the apparent decline. Our results show that rising temperatures combined with linear estimates based on calendar time produce observations of declining sensitivity—without any shift in the underlying biology. Current methods may thus undermine efforts to identify when and how warming will reshape biological processes.

**Significance statement:** Recently a growing body of literature has observed declining temperature sensitivities of plant leafout and other events with higher temperatures. Such results suggest that climate change is already reshaping fundamental biological processes. These temperature sensitivities are often estimated as the magnitude of a biological response per °C from linear regression. The underlying model for many events—that a critical threshold of warmth must be reached to trigger the event—however, is non-linear. We show that this mismatch between the statistical and biological models can produce the illusion of declining sensitivities with warming using current methods. We suggest simple alternative approaches that can better identify when and how warming will reshape biological processes.

## 1 Main text

Climate change has reshaped biological processes around the globe, with shifts in the timing of major life history events (phenology), carbon dynamics and other ecosystem processes (IPCC,2014). With rising temperatures, a growing body of literature has documented changes in temperature sensitivity—the magnitude of a biological response scaled per °C. Many studies have found declining responses to temperature in recent decades (Fu et al., 2015; Güsewell et al.,2017; Piao et al., 2017; Dai et al., 2019) or lower sensitivities in warmer, urban areas (Meng et al., 2020).

Most studies attribute changes in temperature sensitivity to shifts in underlying biological processes. For example, researchers have suggested weaker temperature sensitivities are evidence of increased light limitation in the tundra (Piao et al., 2017), or a decline in the relative importance of warm spring temperatures for spring phenological events (e.g., leafout, insect emergence) in the temperate zone (Fu et al., 2015; Meng et al., 2020), as other environmental triggers (e.g., winter temperatures that determine ‘chilling’) play a larger role. Yet, despite an increase in studies reporting declining or shifting temperature sensitivities, none have provided strong evidence of the biological mechanisms underlying these changes (e.g., Fu et al., 2015; Meng et al.,2020). The missing mechanisms may be hidden in the data: environmental factors moderate biological processes in complex ways (Chuine et al., 2016; Güsewell et al., 2017), are strongly correlated in nature (e.g., Fu et al., 2015), and temperature variance shifts over time and space (Keenan et al., 2020).

Here we propose a simpler alternative explanation: the use of linear models for non-linear responses to temperature. Researchers generally use methods with assumptions of linearity to calculate temperature sensitivities, often relying on some form of linear regression to compute a change in a quantity—days to leafout or carbon sequestered over a fixed time, for example— per °C, thus ignoring that many biological responses to temperature are non-linear. We show, theoretically then with simulated and empirical data, how the use of linear methods for nonlinear responses can produce an illusion that the mechanisms underlying biological processes are changing.

Many observed biological responses are the result of continuous non-linear processes that depend on temperature, which are discretized into temporal units for measurement. For example, a biological response, such as leafout, occurs when a certain thermal sum is reached (Dijkhuis, 1956;Lindsey and Newman, 1956), and plants will reach this threshold more quickly—in calendar time—when average daily temperatures are warmer (Valentine, 1983; Lechowicz, 1984; Kramer,2012). Biologically, however, the plants may require the same temperature sum. Indeed any process observed or measured as the time until reaching a threshold is inversely proportional to the speed at which that threshold is approached. Temperature determines the speed of many biological processes (Bonan and Sirois, 1992; Hinrichsen, 2009; Hofmann and Todgham, 2010). Thus, at very low temperatures plants would never leaf out and at higher temperatures they could leaf out in only a matter of days—yet sensitivities estimated from linear regression at higher (warmer) temperatures would appear much lower than those observed at lower temperatures. Warming acts to step on the biological accelerator, producing shifts in estimates when non-linear responses are modeled as linear.

We show this by deriving the relationship between a biological response and temperature using a simple stochastic model, which describes the first time a random process hits a threshold (see ‘A first-hitting-time model of leafout’ in Supplementary Information). Our model holds the temperature threshold for leafout constant (Hunter and Lechowicz, 1992; Zohner et al., 2020). Even though the mechanism by which temperature leads to leafout does not change, the model produces declining sensitivity—as measured in days per °C—with warming. Indeed, under this model constant temperature sensitivity would be evidence that the temperature threshold is not constant and the mechanisms underlying the leafout process have changed.

Simulations show that correcting for non-linearity using the transformation for an inverse relationship— log transformation—removes apparent declines in temperature sensitivity (Fig. 1, S2, code link). In empirical long-term leafout data from Europe, correcting for non-linearity in responses produces little evidence for declining sensitivities with warming (Figs. 1, S6, S7). An apparent decline in sensitivity for silver birch (*Betula pendula*) from −4.3 days/°C to −3.6 days/°C from 1950-1960 compared to 2000-2010 disappears using a log-log regression (−0.17 versus −0.22). We see similar corrections using 20-year windows, and a potential increase in sensitivity for European beech (*Fagus sylvatica*, see Tables S1-S2). Moreover, the variance of the leafout dates of both species declines as temperatures rise—(declines of roughly 50%, see Tables S1-S2), which is expected under our model as warming accelerates towards the thermal threshold that triggers leafout (and in contrast to predictions from changing mechanisms, see Ford et al., 2016).

**Figure 1:**
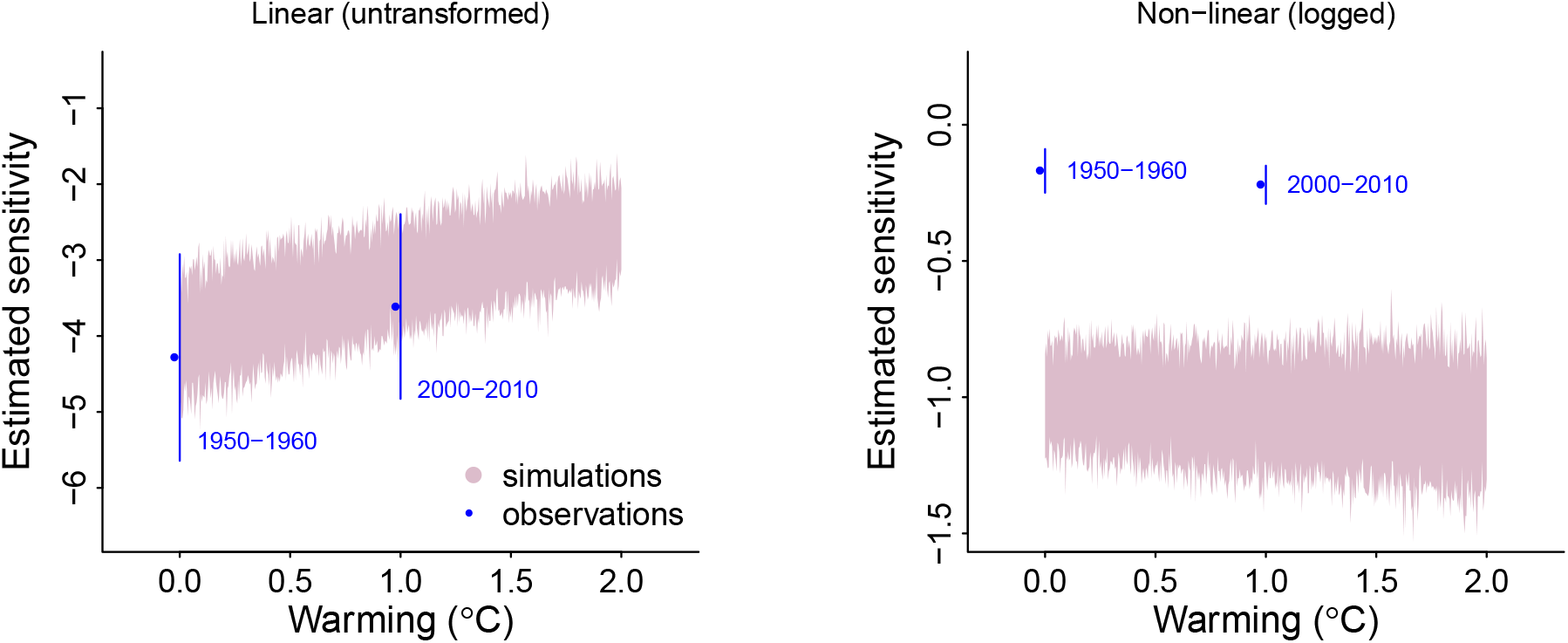
Shifts in temperature sensitivities (response per °C) with warming occur when using linear models for non-linear processes. Estimated sensitivities decline (in magnitude) with warming in simulations (shading, estimated across 45 sites with a base temperature of normal(6,4), variation comes from fluctuation in the Monte Carlo simulations) with no underlying change in the biological process when sensitivities were estimated with linear regression (left). This decline disappears when performing the regression on logged predictor and response variables (right). Such issues may underlie declining sensitivities calculated from observational data, including long-term observations of leafout across Europe (for *Betula pendula* from PEP725 from for the 45 sites that had complete data for 1950-1960 and 2000-2010), which show a lower sensitivity with warming when calculated on raw data, but no change in sensitivity using logged data. Shading, symbols and lines represent means ± standard deviations of regressions across sites. See Supplementary Information for a discussion of why estimated sensitivities are −1 or lower in non-linear models.

Fundamentally rising temperature should alter many biological processes, making robust methods for identifying these changes critical. In spring plant phenology, where declining sensitivities are often reported (Fu et al., 2015; Piao et al., 2017; Dai et al., 2019), warming may increase the role of ‘chilling’ (determined mainly by winter temperatures) and daylength (Laube et al.,2014; Zohner et al., 2016)—potentially increasing the thermal sum required for leafout at lower values of these cues (Polgar et al., 2014; Zohner et al., 2017; Flynn and Wolkovich, 2018). Adjusting our simulations to match this model yielded shifts in sensitivities with warming. Unlike a model with no underlying biological change, however, after correcting for non-linearity, the shifts in sensitivities remained and they occurred in step with the biological change (Fig. S4a, c). In contrast, sensitivities estimated from a linear model showed shifts across the entire range of warming, well before the simulated biological change (Fig. S4a, c). Further, we found that an increase in the thermal sum required for leafout should yield larger in magnitude temperature sensitivities, not smaller, as is often expected (e.g., Fu et al., 2015), thus highlighting the complexity of identifying what trends to expect in sensitivities with warming (see ‘Simulations of common hypotheses for declining sensitivity’ in Supplementary Information for an extended discussion).

Our theoretical model and empirical results show that rising temperatures are sufficient to explain declining temperature sensitivity. It is not necessary to invoke changes to the mechanisms that underlie the biological processes themselves. Our results provide a simpler explanation for observations of declining temperature sensitivities, but do not rule out that important changes in biological processes may underlie such declines. Instead, our results highlight how the use of linear models may make identifying when—and why—warming alters underlying biology far more difficult.

Inferring biological processes from statistical artifacts is not a new problem (e.g., Nee et al.,2005), but climate change provides a new challenge in discerning mechanism from measurements because it affects biological time, while researchers continue to use calendar time. Other fields focused on temperature sensitivity often use approaches that acknowledge the non-linearity of responses (e.g., Yuste et al., 2004). Researchers have called for greater use of process-based models (Keenan et al., 2020), which often include non-linear responses to temperature, but rely themselves on exploratory methods and descriptive analyses for progress (Chuine et al.,2016). The challenge, then, is to interrogate the implicit and explicit models we use to interpret data summaries, and to develop null expectations that apply across biological and calendar time.

## Acknowledgements

Thanks to TJ Davies, A Donnelly, TM Giants, Y. Fu, D. Lipson, C. Rollinson, Y. Vitasse for comments that improved the manuscript. IM-C acknowledges funding from the Spanish Ministry for Science and Innovation. NSERC (grant no. RGPIN05038 to EMW), Canada Research Chair in Temporal Ecology (EMW) and the Spanish Ministry for Science and Innovation (grant no. PID2019/109711RJ-I00 to IM-C) provided funding.

## Data & Code Availability

Code for simulations, PEP 725 analysis, and plots is provided here. For empirical examples, we used PEP 725 phenological data and E-OBS climate data, both of which are freely available via the links.

## List of Supplementary Information

A first-hitting-time model of leafout

Simulations of common hypotheses for declining sensitivity

Methods & results using long-term empirical data (PEP725)

Table S1-S2

Fig S1-S7

