## Supplementary material for "A simple explanation for declining temperature sensitivity with warming": Supp Info

#### 1 A first-hitting-time model of leafout

We use a simple model based on the general understanding of how warm temperatures (forcing) trigger leafout in temperate deciduous trees (Chuine, 2000): that leafout occurs after a certain thermal sum is met. Though very simple, we formalize it here to show its inherent non-linearity and some of its statistical properties.

We use a first-hitting-time model, which describes the first time a random process hits a threshold, because of its broad applicability and conceptual simplicity. We define leafout day, $n_\beta$, as the day, $n$, that cumulative daily temperature, $S^n$, hits the threshold, $\beta$.

We derive the relationship between daily temperature and leafout in two common scenarios. In the first, we take the average daily temperature up until the leafout date. In the second, we take the average daily temperature over a fixed window, such as March 1st to April 30th. In both cases, we discretize time since, although many biological processes depend continuously on time, research typically measures time in discretized units, such as days, weeks, or months.

##### 1.1 Scenario 1: Using average daily temperature until the leafout date

We use the following notation:

$$\begin{aligned}
n &= \text{day since temperatures start to accumulate, } n = 0, 1, ..., N \\
X_n &= \text{observed temperature on day } n \\
S_0^n &= \sum_{i=0}^{n} X_i, \text{the cumulative daily temperature from day 0 to day } n \\
M_0^n &= \frac{S_0^n}{n}, \text{the average daily temperature from day 0 to day } n \\
\beta &= \text{the threshold of interest, } \beta > 0, \text{ (thermal sum required for leafout)} \\
n_\beta &= \underset{n}{\operatorname{argmin}}\ S_n > \beta, \text{the first day the cumulative daily temperature passes the threshold} \\
&\quad \text{(for example, day of year (doy) of leafout).}
\end{aligned}$$

We model $X_n$ as a Gaussian random walk, $X_n \overset{\text{i.i.d}}{\sim} \text{normal}\,(\alpha_0 + \alpha_1 n, \sigma)$, where $\alpha_0 > 0$ is the average temperature on day $n = 0$, $\alpha_1 > 0$ is the day-over-day increase in average temperatures, and $\sigma$ is the standard deviation. This model differs from the traditional Gaussian random walk because of the factor $n$.

This model has two important consequences:

(1) Leafout time is inversely related to average temperature at leafout time.

Under this model, $M_0^{n_\beta}$ and $n_\beta$ are inversely proportional. To see why, assume for the moment that the cumulative daily temperature hits the threshold exactly on leafout day. That is, $S_0^{n_\beta} = \beta$. Then

$$M_0^{n_\beta} = \frac{S_0^{n_\beta}}{n_\beta} = \frac{\beta}{n_\beta}$$

rearranging yields

$$n_\beta = \frac{\beta}{M_0^{n_\beta}}$$

Many global change biology studies use linear regression to quantify the relationship between $n_\beta$ and $M_0^{n_\beta}$ (or similar metrics, see Wolkovich et al., 2012; Piao et al., 2017; Keenan et al., 2020, for examples). Regressing $n_\beta$ on $M_0^{n_\beta}$ finds a best fit line to the inverse curve, $n_\beta = \frac{\beta}{M_0^{n_\beta}}$. The relationship is linearized with the logarithm transformation: $\log(n_\beta) = \log(\beta) - \log(M_0^{n_\beta})$. That is, $\log(n_\beta)$ is linear in log-average daily temperature with slope -1 and intercept $\log(\beta)$.

(2) The variance of the average temperature may decreases as temperatures rise.

Under the model, the mean and variance of $M_0^n$ are $\mathrm{E}(M_0^n|\alpha_0, \alpha_1) = \frac{1}{n}\sum_{i=0}^{n}(\alpha_0 + \alpha_1 i) = \alpha_0 + \alpha_1\frac{(n+1)}{2}$ and $\mathrm{Var}(M_0^n|\alpha_0, \alpha_1) = \frac{\sigma^2}{n}$.

By the law of total variance,

$$\begin{aligned}
\mathrm{Var}(M_0^n) &= \mathrm{E}(\mathrm{Var}(M_0^n|\alpha_0, \alpha_1)) + \mathrm{Var}(\mathrm{E}(M_0^n|\alpha_0, \alpha_1)) \\
&= \frac{\sigma^2}{n} + \mathrm{Var}(\alpha_0 + \alpha_1\frac{n+1}{2}) \\
&= \frac{\sigma^2}{n} + \mathrm{Var}(\alpha_0) + \frac{(n+1)^2}{4}\mathrm{Var}(\alpha_1) + (n+1)\mathrm{Cov}(\alpha_0, \alpha_1)
\end{aligned}$$

As temperatures rise and leafout date becomes earlier, the variance of the average temperature will decline—provided the variation in temperatures, $\sigma^2$, is sufficiently small.

##### 1.2 Scenario 2: Using average daily temperature over a fixed window

We slightly modify the notation:

$$
\begin{aligned}
n &= \text{day since temperatures start to accumulate, } n = 0, \ldots, a, \ldots, b \\
X_n &= \text{observed temperature on day } n \\
S_a^n &= \sum_{i=a}^{n} X_i, \text{the cumulative daily temperature from day } a \text{ to day } n \\
M_a^n &= \frac{S_a^n}{n-a}, \text{the average daily temperature from day } a \text{ to day } n \\
\beta &= \text{the threshold of interest, } \beta > 0, \text{ (thermal sum required for leafout)} \\
n_\beta &= \underset{n}{\text{argmin}}\ S_0^n > \beta, \text{the first day the cumulative daily temperature passes the threshold} \\
&\quad \text{(for example, day of year (doy) of leafout).}
\end{aligned}
$$

As before, we model $X_n$ as a Gaussian random walk, $X_n \overset{\text{i.i.d}}{\sim} \text{normal}\left(\alpha_0 + \alpha_1 n, \sigma\right)$, where $\alpha_0 > 0$ is the average temperature on day $n = 0$, $\alpha_1 > 0$ is the day-over-day increase in average temperatures, and $\sigma$ is the standard deviation. We make the additional assumption that $X_n \geq 0$ for all $n$ and $a < n_\beta < b$. That is, the cumulative temperature acquired by the plant always increases.

Note that

$$S_a^b \sim \text{normal}\left(\alpha_0(b-a) + \tfrac{\alpha_1}{2}(b-a)(b+a+1), \sigma\sqrt{b-a}\right)$$

$$M_a^b \sim \text{normal}\left(\alpha_0 + \tfrac{\alpha_1}{2}(b+a+1), \tfrac{\sigma}{\sqrt{b-a}}\right)$$

$$S_n^b - S_a^b \sim \text{normal}\left(\alpha_0(b-a-n) + \tfrac{\alpha_1}{2}((b-n)(b+n+1) - a(a+1)), \sigma\sqrt{b+a-n}\right)$$

so that

$$
\begin{aligned}
Pr\left(n_\beta \leq n \mid M_a^b = m\right) &= Pr\left(n_\beta \leq n \mid S_a^b = (b-a)m\right) \\
&= Pr\left(S_0^n \geq \beta \mid S_a^b = (b-a)m\right) \\
&= Pr\left(S_n^b \leq (b-a)m + S_0^a - \beta\right) \\
&= Pr\left(S_n^b - S_0^a \leq (b-a)m - \beta\right) \\
&= \Phi\left(\frac{(b-a)m - \beta - [\alpha_0(b-a-n) + \frac{\alpha_1}{2}((b-n)(b+n+1) - a(a+1))]}{\sigma\sqrt{b+a-n}}\right)
\end{aligned}
$$

The distribution of $M_a^b$ shows that consequence (2) above still holds with this model. Consequence (1) no longer holds directly, but will in many situations where average daily temperature until an event correlates strongly with average daily temperature because the window is chosen based, in part, on the expected hitting time (Figs. S1-S2). We note two additional consequences:

(3) The conditional median is quadratic in $n$:

$$
\begin{aligned}
\frac{1}{2} &\overset{\text{set}}{=} Pr\left(n_\beta \leq n \mid M_a^b = m\right) \\
&\Rightarrow 0 = (b-a)m - \beta - [\alpha_0(b-a-n) + \frac{\alpha_1}{2}((b-n)(b+n+1) - a(a+1))] \\
&\Rightarrow m = \frac{1}{(a-b)}[-\beta - \alpha_0(b-a-n) - \frac{\alpha_1}{2}((b-n)(b+n+1) - a(a+1))] \\
&\quad = \frac{1}{(a-b)}[-\beta - \alpha_0(b-a) - \frac{\alpha_1}{2}(b-a)(b+a+1)] + \frac{\alpha_0 + \frac{\alpha_1}{2}}{(a-b)}n + \frac{\frac{\alpha_1}{2}}{(a-b)}n^2 \\
&\quad := \gamma_0 + \gamma_1 n + \gamma_2 n^2
\end{aligned}
$$

(4) The conditional mean and variance are sums of negative sigmoids, according to the following identities

$$E\left(n_\beta \mid M_a^b = m\right) = \sum_{n=0}^{\infty} Pr\left(n_\beta \geq n \mid M_a^b = m\right)$$

$$E\left(n_\beta^2 \mid M_a^b = m\right) = \sum_{n=0}^{\infty} n\ Pr\left(n_\beta \geq n \mid M_a^b = m\right)$$

### 2 Simulations of common hypotheses for declining sensitivity

#### 2.1 Effect of increasing thermal sum on sensitivity

Many biological hypotheses explain the observed decline in plant sensitivity as a consequence of declines in over-winter chilling or short photoperiods (Fu et al., 2015; Zohner et al., 2016; Fu et al., 2019). These forces generally increase the thermal sum required for leafout each year (Polgar et al., 2014; Zohner et al., 2017; Flynn and Wolkovich, 2018), hypothesized to weaken the linear relationship between biological responses and temperature.

When the thermal sum required for leafout ($\beta$ in our first hitting-time model in section 1.1 above) increases the regression coefficient from regressing leafout date on spring-time temperature increases in magnitude, that is, it leads to increases (not declines) in sensitivity (Fig. S3). This happens due to a decreasing (in magnitude) variance in temperature alongside an increasing in magnitude covariance between temperature and leafout day. Similar results are seen given the model in section 1.2, though due mainly to increasing covariance between temperature and leafout day.

#### 2.2 Effects of chilling and daylength

We extended simulations from our first hitting-time model to examine two common biological hypotheses for declining sensitivities in spring plant phenology to understand how discernible biological shifts would be from the shifts present when using linear models to estimate sensitivities for non-linear temperature responses.

First, we simulated the most often cited hypothesis (e.g., Fu et al., 2015; Piao et al., 2017): an increasing thermal sum threshold given declines in over-winter chilling with warming. Experiments in controlled environments show that plants require greater thermal sums to budburst or leafout given lower chilling (e.g., Laube et al., 2014; Flynn and Wolkovich, 2018). Connecting chilling to leafout in observational data, however, is difficult as we know little mechanistically about what temperatures determine chilling (but see Rinne et al., 2011; van der Schoot et al., 2014, for recent developments in lab molecular work), and most models of chilling are based on statistical correlations (e.g., Erez and Lavee, 1971; Richardson, 1974; Luedeling et al., 2009).

Without robust estimates of chilling, teasing out effects of chilling from forcing is difficult. Necessarily detailed models are non-identified (data for a particular model are non-identifiable when multiple parameter(s) values can be estimated from the same data). In the example of our simulated chilling + forcing model, the same leafout day can be driven by a lower thermal sum ($\beta$ in our first-hitting time model) when chilling is higher, and higher thermal sums when chilling is lower, making chilling and forcing parameter values both non-identifiable. The model could become identifiable with an accurate covariate predicting chilling or forcing, which underlies efforts to model chilling (Luedeling and Brown, 2011; Harrington and Gould, 2015) and 'pre-season' length (Güsewell et al., 2017; Xu et al., 2018, and discussed below in 'Methods & results using long-term empirical data (PEP725)'), but these efforts are mostly based on statistical fits, which cannot easily overcome non-identifiability without additional information.

In a model where increased thermal sums are driven by declines in overwinter chilling with warming sensitivities from log-transformed data first increase and then decrease (as winter warming delays leafout day and spring warming advances leafout day) in step with biological shifts in cues (Fig. S4a, c), while for estimates from the linear model changes in sensitivity occur throughout warming, despite no major change in cues before 4 °C (Fig. S4a, c). The ultimate effect with warming depends on exact parameter values (e.g., how much thermal cues shift with declines in warming) and likely on the covariation of $X$ and $\beta$, highlighting how difficult teasing apart relationships in observational spring phenology may be given correlations across multiple predictors (correlations that are likely exacerbated by climate change).

Second, we simulated an alternative hypothesis where warming causes the thermal sum to be reached before a required daylength threshold and plants then leaf out on the first day the daylength threshold is met (Zohner et al., 2016; Fu et al., 2019). At its extreme this model produces the same leafout day (the day when the required daylength is met) across different temperatures and thus produces smaller in magnitude estimated sensitivities in both linear and logged models. Estimated sensitivities, however, from a linear model do not necessarily decline depending on exact parameter values (see Fig. S4b, d). Again this shift appears more visible when using sensitivities that include the non-linear nature of plants temperature response compared to linear estimates (see Fig. S4b, d).

Annotated code for these simulations are available via github to allow testing of alternative parameter values or versions of these hypotheses.

#### 3 Methods & results using long-term empirical data (PEP725)

To examine how estimated sensitivities shift over time, we selected sites of two common European tree species (silver birch, *Betula pendula*, and European beech, *Fagus sylvatica*) that have long-term observational data of leafout, through the Pan European Phenology Project (PEP725, Templ et al., 2018). We selected these two species given that they were best represented for consistent data at the same sites over our study years for an early-leafout (*Betula pendula*) and a late-leafout (*Fagus sylvatica*) species (e.g., *Betula pendula* had 17 sites with leafout data from 1950-1960 and 2000-2010, while the next best option for an early-leafout species, *Alnus glutinosa*, had data for only five sites). We used sites with complete leafout data across both our 10-year (and 20-year) windows to avoid possible confounding effects of shifting sites over time (see Tables S1-S2 for numbers of sites per species-window combination).

To calculate temperature sensitivities, we used a European-wide gridded climate dataset *(*E-OBS*, Cornes et al., 2018)* to extract daily minimum and maximum temperature for the grid cells where observations of leafout for these two species were available (Fig. S5 shows a subset of the climate data for 14 sites used). Determining the appropriate window over which to estimate a temperature sensitivity for spring plant phenology is an area of active research (Güsewell et al., 2017; Xu et al., 2018). Ideally researchers wish to separate windows over which chilling and forcing apply (if they can be cleanly separated, Linkosalo et al., 2008; Lundell et al., 2020), but this is generally impossible given our limited understanding of the two processes (Chuine et al., 2016). Researchers thus either use a pre-defined spring window (e.g., Park et al., 2019; Kopp et al., 2020) or use a statistical search to determine window attributes; for example, some use a set period then search for a start date (e.g., Cook et al., 2012), while others search for both a start date and window length (e.g., Fu et al., 2015; Tansey et al., 2017). Given the non-identifiability of the simple chilling + forcing model described above ('Simulations of common hypotheses for declining sensitivity'), we do not feel there is sufficient evidence that statistical searches for pre-season windows will select biologically relevant periods related to forcing, and more easily could add additional layers of non-identifiability (to date, most pre-season windows are fit as separate analyses making such issues harder for researchers to detect). Thus, we used pre-determined windows from 1 March to 30 April (60 d; we also present windows of 45 d, from 1 Mar to 15 Apr, and for 31 d, from 15 Mar to 15 Apr, for comparison). Our window represents a period in the spring when forcing is likely the dominant cue, and is similar to many other studies of temperature plants using pre-defined windows (e.g., Bolmgren et al., 2013; Prevey et al., 2017; Park et al., 2019). We then estimated the sensitivity as a simple linear regression of leafout day of year versus the mean daily temperature over the window ('slope' in Tables S1-S2), or a simple linear regression using logged versions of these predictors ('log-slope' in Tables S1-S2). Code for these analyses are available via github (`https://github.com/temporalecologylab/labgit/tree/master/projects/decsenspost`).

Our estimates of temperature sensitivity from a linear model using untransformed variables show a decline in sensitivity with recent warming for *Betula pendula* over 10 and 20-year windows, but no decline for *Fagus sylvatica*; using logged variables estimates appeared more similar over time or sometimes suggested an increase in sensitivity (see Figs. S6-S7, Tables S1-S2). This shift in estimated sensitivity when regressing with untransformed versus logged variables suggests the declining estimates with untransformed variables may not be caused by changes in the underlying mechanisms of leafout (i.e., reduced winter chilling) and driven instead by using linear regression for a non-linear process. This hypothesis is supported further by large declines in variance of leafout in recent decades.

Shifts in variance provide another hurdle to robust estimates of temperature sensitivity. Previous work has highlighted how shifting temperature variance (over space and/or time) could lead to shifting estimates of temperature sensitivities (Keenan et al., 2020), but our results stress that variance in both leafout and temperature are shifting. If both shift in step, estimates would not be impacted by changes in temperature variance, but our results suggest variance in temperature—for these data—has declined more than variance in leafout, though both have declined substantially in recent decades (Tables S1-S2).

Estimated sensitivities for the empirical data (PEP725) using logged variables are far lower than the value obtained in our simulations (-1). This likely results from a contrast between our simulations—where we can accurately define the temperature plants experience and the temporal window that drives leafout—and our empirical data, where we do not know how measured temperatures translate into the temperatures that plants accumulate and where we have no clear method to define the relevant temporal window (discussed above).

These results highlight how the acceleration of biological time due to climate change requires researchers to clarify their assumptions. Expecting temperature sensitivity to remain constant as temperatures rise assumes the relationship between response and temperature is proportional. But the underlying biological processes suggest this relationship is seldom proportional, or even linear. In fact, when our model holds, declining sensitivity with rising temperatures should be the null hypothesis of any analysis of temperature sensitivity based on linear regression or similar methods.

*References*

Bolmgren, K., D. Vanhoenacker, and A. J. Miller-Rushing. 2013. One man, 73 years, and 25 species. evaluating phenological responses using a lifelong study of first flowering dates. International Journal of Biometeorology 57:367–375.

Chuine, I. 2000. A unified model for budburst of trees. Journal of Theoretical Biology 207:337–347.

Chuine, I., M. Bonhomme, J.-M. Legave, I. García de Cortázar-Atauri, G. Charrier, A. Lacointe, and T. Améglio. 2016. Can phenological models predict tree phenology accurately in the future? The unrevealed hurdle of endodormancy break. Global Change Biology 22:3444–3460.

Cook, B. I., E. M. Wolkovich, and C. Parmesan. 2012. Divergent responses to spring and winter warming drive community level flowering trends. Proceedings of the National Academy of Sciences of the United States of America 109:9000–9005. Cook, Benjamin I. Wolkovich, Elizabeth M. Parmesan, Camille.

Cornes, R. C., G. van der Schrier, E. J. van den Besselaar, and P. D. Jones. 2018. An ensemble version of the E-OBS temperature and precipitation data sets. Journal of Geophysical Research: Atmospheres 123:9391–9409.

Erez, A., and S. Lavee. 1971. Effect of climatic conditions on dormancy development of peach buds. i. temperature. Amer Soc Hort Sci J .

Flynn, D. F. B., and E. M. Wolkovich. 2018. Temperature and photoperiod drive spring phenology across all species in a temperate forest community. New Phytologist 219:1353–1362.

Fu, Y. H., S. Piao, X. Zhou, X. Geng, F. Hao, Y. Vitasse, and I. A. Janssens. 2019. Short photoperiod reduces the temperature sensitivity of leaf-out in saplings of fagus sylvatica but not in horse chestnut. Global change biology 25:1696–1703.

Fu, Y. S. H., H. F. Zhao, S. L. Piao, M. Peaucelle, S. S. Peng, G. Y. Zhou, P. Ciais, M. T. Huang, A. Menzel, J. P. Uelas, Y. Song, Y. Vitasse, Z. Z. Zeng, and I. A. Janssens. 2015. Declining global warming effects on the phenology of spring leaf unfolding. Nature 526:104–107.

Güsewell, S., R. Furrer, R. Gehrig, and B. Pietragalla. 2017. Changes in temperature sensitivity of spring phenology with recent climate warming in Switzerland are related to shifts of the preseason. Global Change Biology 23:5189–5202.

Harrington, C. A., and P. J. Gould. 2015. Tradeoffs between chilling and forcing in satisfying dormancy requirements for pacific northwest tree species. Frontiers in Plant Science 6:120.

Keenan, T. F., A. D. Richardson, and K. Hufkens. 2020. On quantifying the apparent temperature sensitivity of plant phenology. New Phytologist 225:1033–1040.

Kopp, C. W., B. M. Neto-Bradley, L. P. J. Lipsen, J. Sandhar, and S. Smith. 2020. Herbarium records indicate variation in bloom-time sensitivity to temperature across a geographically diverse region. International Journal of Biometeorology 64:873–880.

Laube, J., T. H. Sparks, N. Estrella, J. Höfler, D. P. Ankerst, and A. Menzel. 2014. Chilling outweighs photoperiod in preventing precocious spring development. Global Change Biology 20:170–182.

Linkosalo, T., H. K. Lappalainen, and P. Hari. 2008. A comparison of phenological models of leaf bud burst and flowering of boreal trees using independent observations. Tree Physiology 28:1873–1882.

Luedeling, E., and P. H. Brown. 2011. A global analysis of the comparability of winter chill models for fruit and nut trees. International Journal of Biometeorology 55:411–421.

Luedeling, E., M. H. Zhang, G. McGranahan, and C. Leslie. 2009. Validation of winter chill models using historic records of walnut phenology. Agricultural and Forest Meteorology 149:1854–1864.

Lundell, R., H. Hanninen, T. Saarinen, H. Astrom, and R. Zhang. 2020. Beyond rest and quiescence (endodormancy and ecodormancy): A novel model for quantifying plant-environment interaction in bud dormancy release. Plant Cell and Environment 43:40–54.

Park, D. S., I. Breckheimer, A. C. Williams, E. Law, A. M. Ellison, and C. C. Davis. 2019. Herbarium specimens reveal substantial and unexpected variation in phenological sensitivity across the eastern united states. Philosophical Transactions of the Royal Society B: Biological Sciences 374:20170394.

Park, H., S. J. Jeong, C. H. Ho, C. E. Park, and J. Kim. 2018. Slowdown of spring green-up advancements in boreal forests. Remote Sensing of Environment 217:191–202.

Piao, S., Z. Liu, T. Wang, S. Peng, P. Ciais, M. Huang, A. Ahlstrom, J. F. Burkhart, F. Chevallier, I. A. Janssens, et al. 2017. Weakening temperature control on the interannual variations of spring carbon uptake across northern lands. Nature Climate Change 7:359.

Polgar, C., A. Gallinat, and R. B. Primack. 2014. Drivers of leaf-out phenology and their implications for species invasions: insights from Thoreau's Concord. New Phytologist 202:106–15.

Prevey, J., M. Vellend, N. Ruger, R. D. Hollister, A. D. Bjorkman, I. H. Myers-Smith, S. C. Elmendorf, K. Clark, E. J. Cooper, B. Elberling, A. M. Fosaa, G. H. R. Henry, T. T. Hoye, I. S. Jonsdottir, K. Klanderud, E. Levesque, M. Mauritz, U. Molau, S. M. Natali, S. F. Oberbauer, Z. A. Panchen, E. Post, S. B. Rumpf, N. M. Schmidt, E. A. G. Schuur, P. R. Semenchuk, T. Troxler, J. M. Welker, and C. Rixen. 2017. Greater temperature sensitivity of plant phenology at colder sites: implications for convergence across northern latitudes. Global Change Biology 23:2660–2671.

Richardson, E. 1974. A model for estimating the completion of rest for 'Redhaven' and 'Elberta' peach trees. HortScience 9:331–332.

Rinne, P. L. H., A. Welling, J. Vahala, L. Ripel, R. Ruonala, J. Kangasjarvi, and C. van der Schoot. 2011. Chilling of dormant buds hyperinduces FLOWERING LOCUS T and recruits GA-Inducible 1,3-beta-Glucanases to reopen signal conduits and release dormancy in *Populus*. Plant Cell 23:130–146.

Tansey, C. J., J. D. Hadfield, and A. B. Phillimore. 2017. Estimating the ability of plants to plastically track temperature-mediated shifts in the spring phenological optimum. Global Change Biology 23:3321–3334.

Templ, B., E. Koch, K. Bolmgren, M. Ungersböck, A. Paul, H. Scheifinger, T. Rutishauser, M. Busto, F.-M. Chmielewski, L. Hájková, S. Hodzić, F. Kaspar, B. Pietragalla, R. Romero-Fresneda, A. Tolvanen, V. Vučetič, K. Zimmermann, and A. Zust. 2018. Pan European Phenological database (PEP725): a single point of access for European data. International Journal of Biometeorology 62:1109–1113.

van der Schoot, C., L. K. Paul, and P. L. H. Rinne. 2014. The embryonic shoot: a lifeline through winter. Journal of Experimental Botany 65:1699–1712.

Wang, C., R. Y. Cao, J. Chen, Y. H. Rao, and Y. H. Tang. 2015. Temperature sensitivity of spring vegetation phenology correlates to within-spring warming speed over the northern hemisphere. Ecological Indicators 50:62–68.

Wolkovich, E. M., B. I. Cook, J. M. Allen, T. M. Crimmins, J. L. Betancourt, S. E. Travers, S. Pau, J. Regetz, T. J. Davies, N. J. B. Kraft, T. R. Ault, K. Bolmgren, S. J. Mazer, G. J. McCabe, B. J. McGill, C. Parmesan, N. Salamin, M. D. Schwartz, and E. E. Cleland. 2012. Warming experiments underpredict plant phenological responses to climate change. Nature 485:494–497.

Xu, Y. J., H. J. Wang, Q. S. Ge, C. Y. Wu, and J. H. Dai. 2018. The strength of flowering-temperature relationship and preseason length affect temperature sensitivity of first flowering date across space. International Journal of Climatology 38:5030–5036.

Zhang, H. C., W. P. Yuan, S. G. Liu, W. J. Dong, and Y. Fu. 2015. Sensitivity of flowering phenology to changing temperature in china. Journal of Geophysical Research-Biogeosciences 120:1658–1665.

Zohner, C. M., B. M. Benito, J. D. Fridley, J. C. Svenning, and S. S. Renner. 2017. Spring predictability explains different leaf-out strategies in the woody floras of North America, Europe and East Asia. Ecology Letters 20:452–460.

Zohner, C. M., B. M. Benito, J. C. Svenning, and S. S. Renner. 2016. Day length unlikely to constrain climate-driven shifts in leaf-out times of northern woody plants. Nature Climate Change 6:1120–1123.

### 4 Tables

Table S1: Climate and phenology statistics for two species (*Betula pendula, Fagus sylvatica*, across 45 and 47 sites respectively) from the PEP725 data across all sites with continuous data from 1950-1960 and 2000-2010. ST is spring temperature from 1 March to 30 April, ST.lo is temperature 30 days before leafout, and GDD is growing degree days 30 days before leafout. Slope represents the estimated sensitivity using untransformed leafout and ST, while log-slope represents the estimated sensitivity using log(leafout) and log(ST). We calculated all metrics for each species x site x 10 year period before taking mean or variance estimates. See also Fig. S6.

| | | mean (ST) | | | | var (ST) | | | | | slope | | | log-slope | | |
|---|---|---|---|---|---|---|---|---|---|---|---|---|---|---|---|---|
| years | species | 31 | 45 | 60 | mean ST.lo | 31 | 45 | 60 | var (lo) | GDD | 31 | 45 | 60 | 31 | 45 | 60 |
| 1950-1960 | *Betula* | 6.3 | 5.2 | 5.6 | 7.0 | 2.0 | 2.6 | 3.4 | 110.5 | 71.7 | 3.3 | -2.1 | -4.3 | 0.20 | -0.09 | -0.17 |
| 2000-2010 | *Betula* | 6.1 | 4.9 | 6.6 | 6.8 | 3.7 | 2.4 | 1.2 | 47.0 | 64.6 | -0.1 | 0.5 | -3.6 | 0.00 | 0.02 | -0.22 |
| 1950-1960 | *Fagus* | 6.3 | 5.3 | 5.6 | 7.5 | 1.9 | 2.6 | 3.3 | 71.9 | 83.8 | 2.0 | -0.9 | -2.8 | 0.12 | -0.05 | -0.11 |
| 2000-2010 | *Fagus* | 6.2 | 5.0 | 6.7 | 7.7 | 3.8 | 2.4 | 1.2 | 38.3 | 86.7 | -0.7 | 1.2 | -3.4 | -0.03 | 0.06 | -0.20 |

Table S2: Climate and phenology statistics for two species (*Betula pendula, Fagus sylvatica*, across 17 and 24 sites respectively) from the PEP725 data across all sites with continuous data from 1950-2010. ST is spring temperature from 1 March to 30 April, ST.lo is temperature 30 days before leafout, and GDD is growing degree days 30 days before leafout. Slope represents the estimated sensitivity using untransformed leafout and ST, while log-slope represents the estimated sensitivity using log(leafout) and log(ST). We calculated all metrics for each species x site x 20 year period before taking mean or variance estimates. See also Fig. S7.

| | | mean (ST) | | | | var (ST) | | | | | slope | | | log-slope | | |
|---|---|---|---|---|---|---|---|---|---|---|---|---|---|---|---|---|
| years | species | 31 | 45 | 60 | mean ST.lo | 31 | 45 | 60 | var (lo) | GDD | 31 | 45 | 60 | 31 | 45 | 60 |
| 1950-1970 | *Betula* | 6.4 | 4.9 | 5.8 | 7.1 | 3.7 | 2.7 | 2.6 | 79.9 | 72.5 | 1.1 | -1.0 | -4.3 | 0.08 | -0.03 | -0.19 |
| 1970-1990 | *Betula* | 6.4 | 5.4 | 5.9 | 7.2 | 2.2 | 2.9 | 1.3 | 104.8 | 72.2 | -0.0 | -2.0 | -6.1 | -0.02 | -0.07 | -0.33 |
| 1990-2010 | *Betula* | 5.8 | 5.3 | 6.8 | 6.7 | 2.1 | 2.7 | 0.9 | 36.2 | 60.0 | -1.2 | 0.0 | -3.3 | -0.07 | 0.00 | -0.21 |
| 1950-1970 | *Fagus* | 6.1 | 4.7 | 5.6 | 7.6 | 3.8 | 2.8 | 2.7 | 63.4 | 86.0 | 1.0 | -0.2 | -3.1 | 0.05 | 0.00 | -0.12 |
| 1970-1990 | *Fagus* | 6.2 | 5.2 | 5.6 | 7.5 | 2.3 | 3.0 | 1.3 | 56.2 | 81.3 | -0.2 | -1.3 | -2.5 | -0.01 | -0.04 | -0.16 |
| 1990-2010 | *Fagus* | 5.5 | 5.2 | 6.7 | 7.5 | 2.2 | 2.8 | 1.0 | 32.8 | 79.9 | -0.6 | 0.1 | -2.8 | -0.03 | 0.01 | -0.15 |

#### 5 Figures

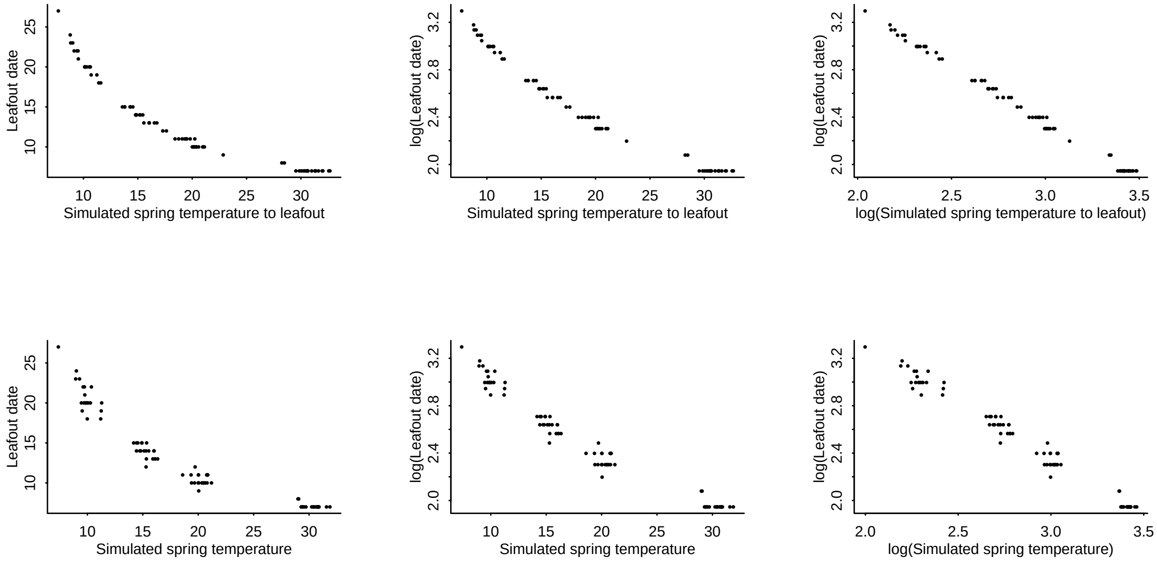


Figure S1: **Simulated leafout as a function of temperature across different temperatures highlights non-linearity of process.** Here we simulated sets of data where leafout constantly occurs at 200 growing degree days (thermal sum of mean daily temperatures with 0°C as base temperature) across mean temperatures of 10, 15, 20 and 30°C (constant SD of 4), we calculated estimated mean temperature until leafout date (top row) or across a fixed window (bottom row, similar to estimates of 'spring temperature'). While within any small temperature range the relationship may appear linear, its non-linear relationship becomes clear across the greater range shown here (left). Taking the log of both leafout and temperature (right) linearizes the relationship.

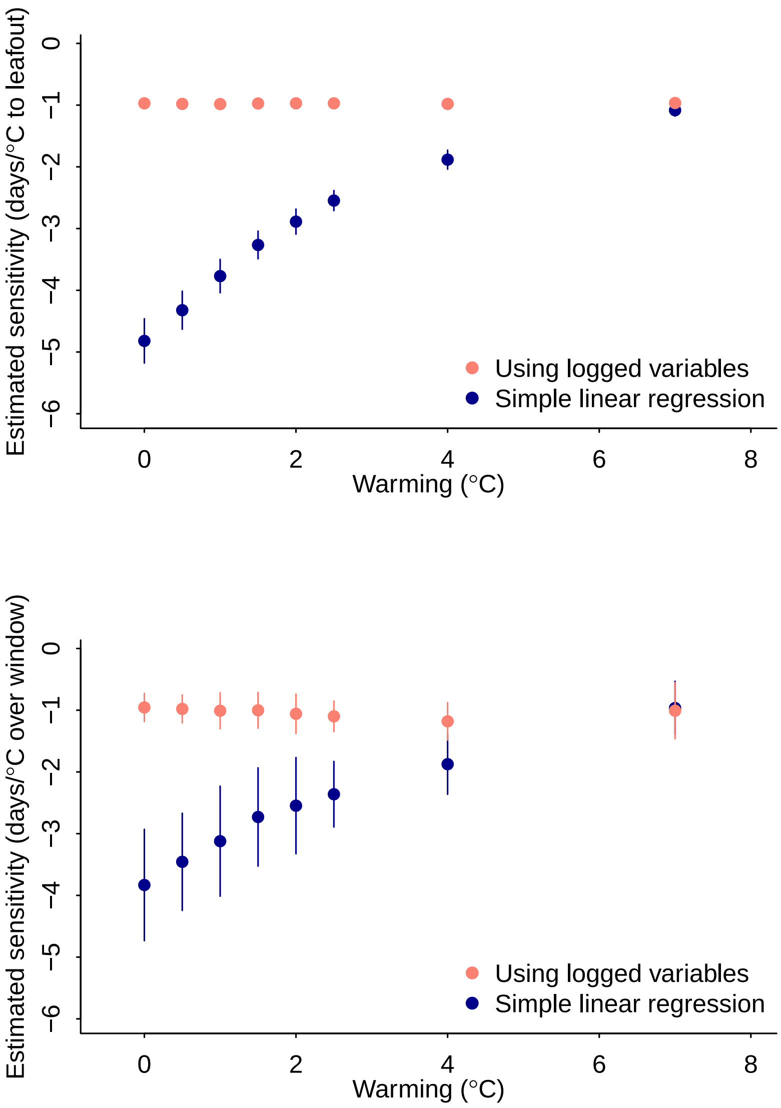


Figure S2: **A simple model generates declining sensitivities with warming.** We show declines in estimated sensitivities with warming from simulations (top: using average temperature until leafout, bottom: using a fixed window; dots and lines represent means ± standard deviation) with no underlying change in the biological process when sensitivities were estimated with simple linear regression ("Simple linear regression"). This decline disappears using regression on logged predictor and response variables ("Using logged variables").

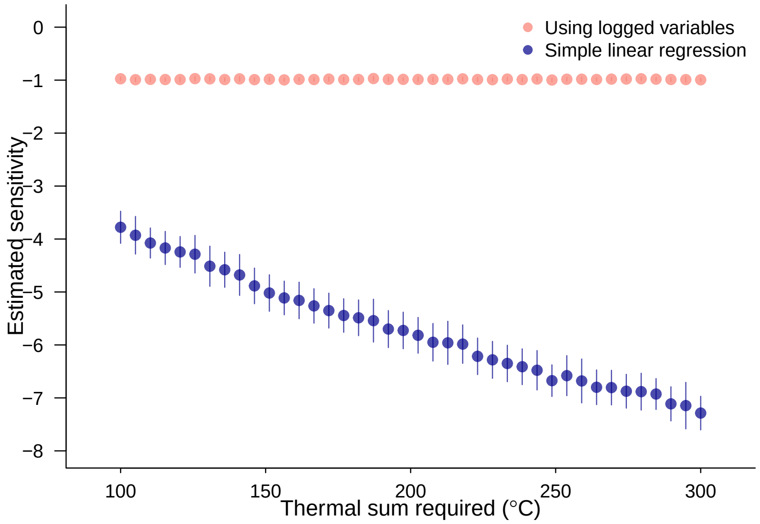


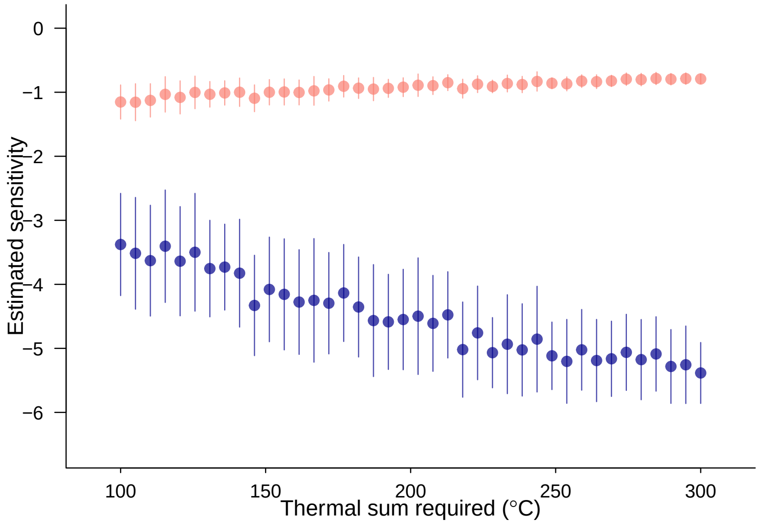


Figure S3: **Simulated leafout as a function of required thermal sum for leafout.** Here we simulated sets of data where leafout occurs at varying thermal sums (sum of mean daily temperatures with 0°C as base temperature) and estimated sensitivities using mean temperature until leafout date (top) or across a fixed window (bottom) with simple linear regression ("Simple linear regression"), and using regression on logged predictor and response variables ("Using logged variables"). Dots and lines represent means ± standard deviation.

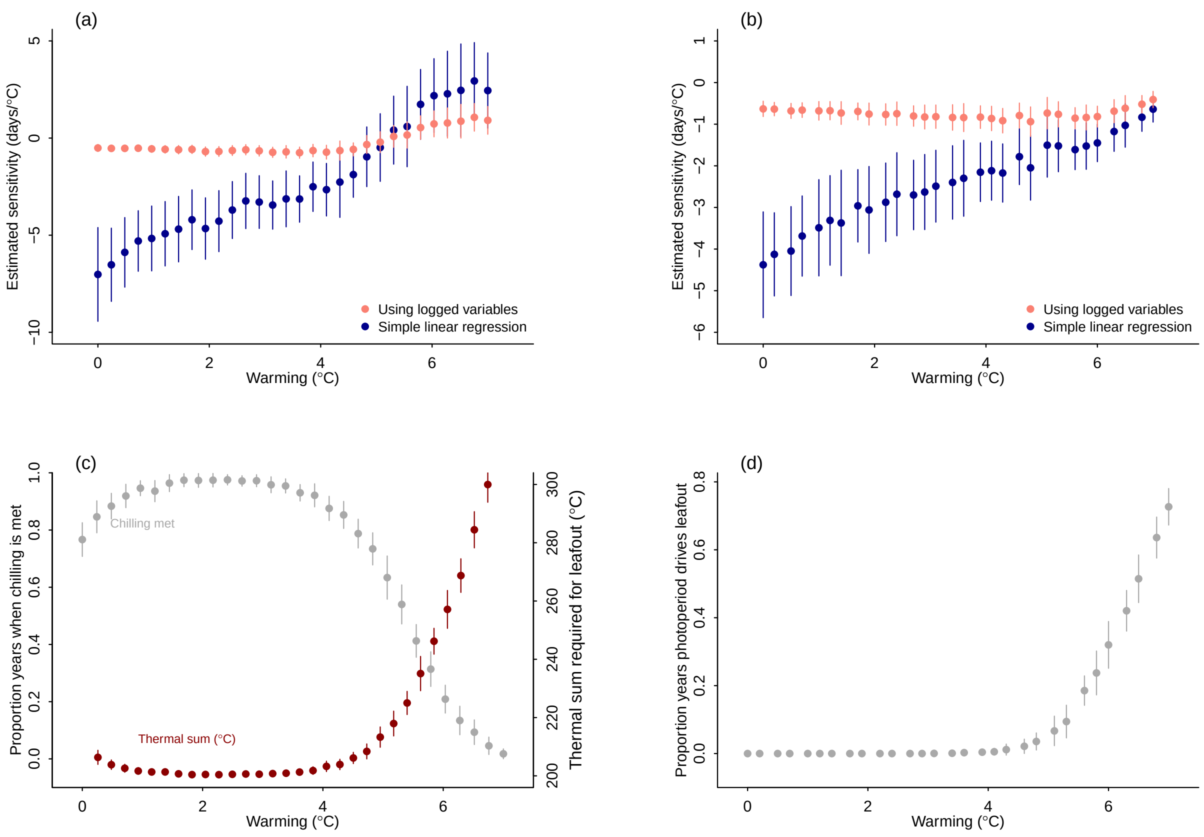


Figure S4: **Simulated leafout as a function of temperature across different levels of warming with shifts in underlying biology through lower chilling (a, c) and photoperiod thresholds (b, d).** We show estimated sensitivities in the top panels (a-b), and the shifting cues in the bottom panels (c-d). Here we simulated sets of data where leafout occurs at a thermal sum of 200 (sum of mean daily temperatures with 0°C as base temperature) when chilling and photoperiod requirements are met, and requires a higher thermal sum when chilling is not met (a, c, required chilling set at 110 units summing temperatures between 0-5°C over 120 d winter period, increasing the thermal sum by 3 units per unmet chilling unit) or where leafout occurs at a thermal sum of 200 as long as the daylength of that day is $\geq$ 12 hours (at 45 N, estimated from R's geosphere package); otherwise leafout occurs on the first day when daylength is 12 hours (b, d). In all simulations the daily increase in spring temperatures was 0.1°C, the variance in daily temperatures was 4°C, in the chilling simulations winter temperatures were centered at 1°C and spring temperatures at 2°C; in the photoperiod simulations spring temperatures were centered at 4°C. Dots and lines represent means $\pm$ standard deviation.

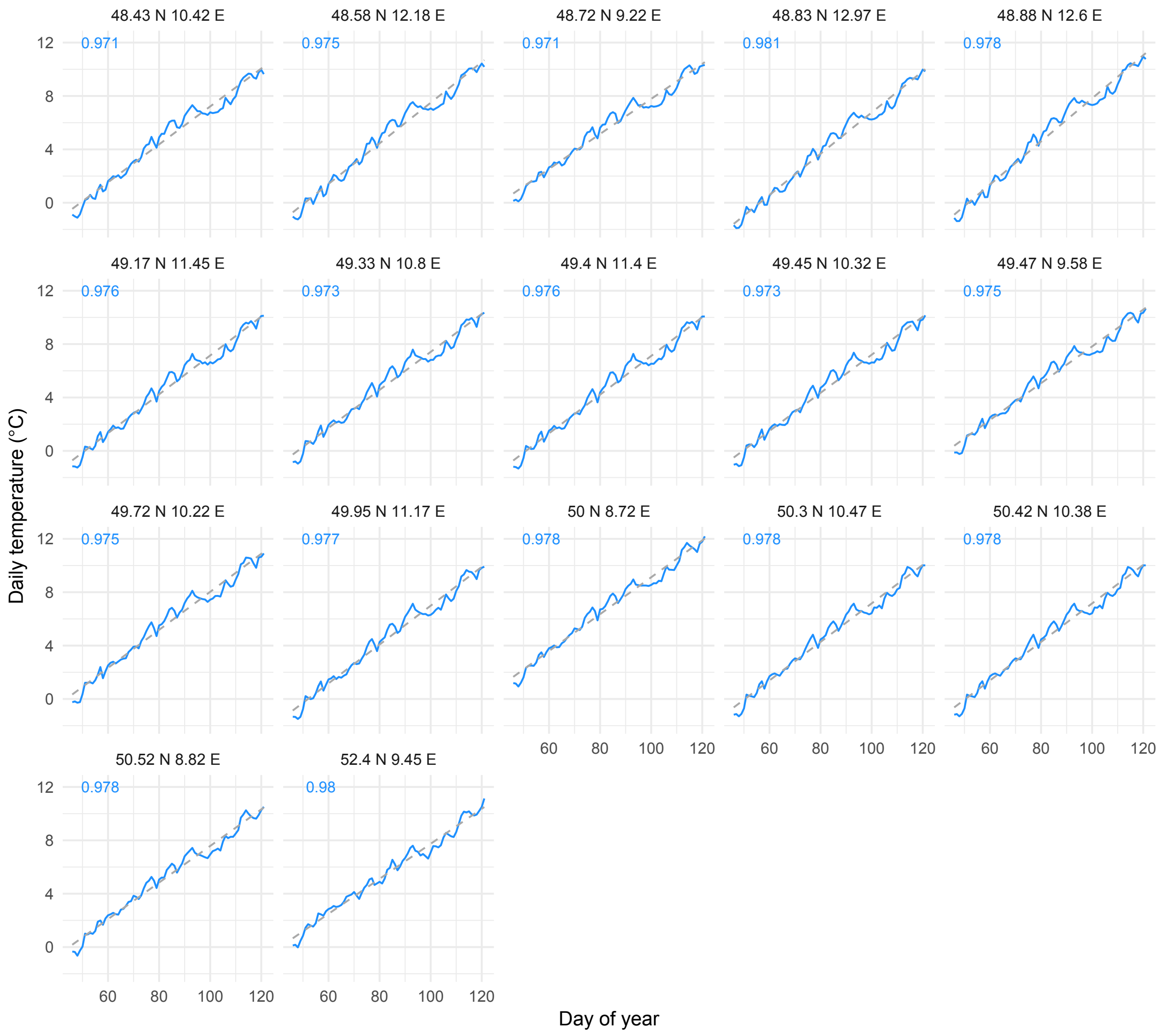


Figure S5: **Mean daily spring temperatures** averaged over 1951-2010 for the 17 PEP725 sites with continuous data from 1950-2010 for *Betula pendula*, with latitude and longitude of each site given on top of each panel; dashed lines show simple linear fits with the $R^2$ given in the light blue for each panel.

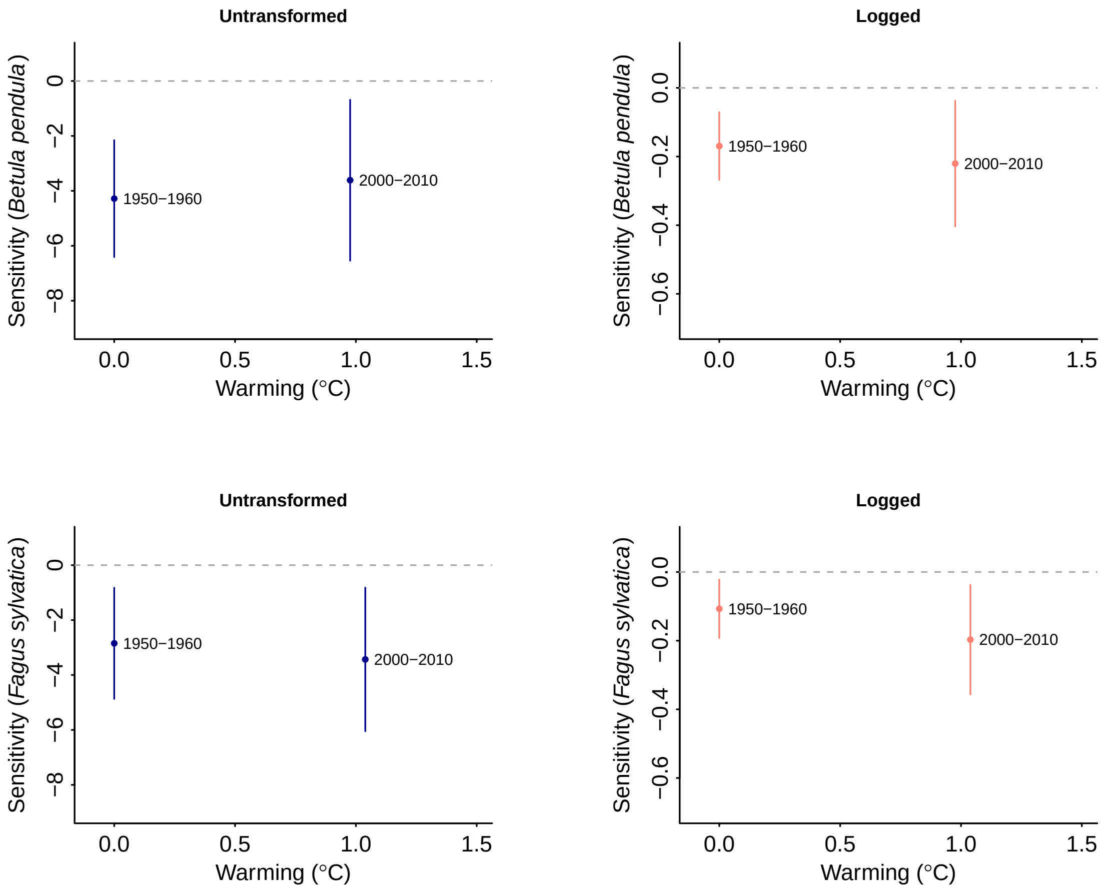


Figure S6: **Sensitivities from PEP725 data using 10 year windows of data** for two species (top – *Betula pendula*, bottom – *Fagus sylvatica*; all lines show 78% confidence intervals from linear regressions). Amounts of warming are calculated relative to 1950-1960 and we used only sites with leafout data in all years shown here. See Table S1 for further details.

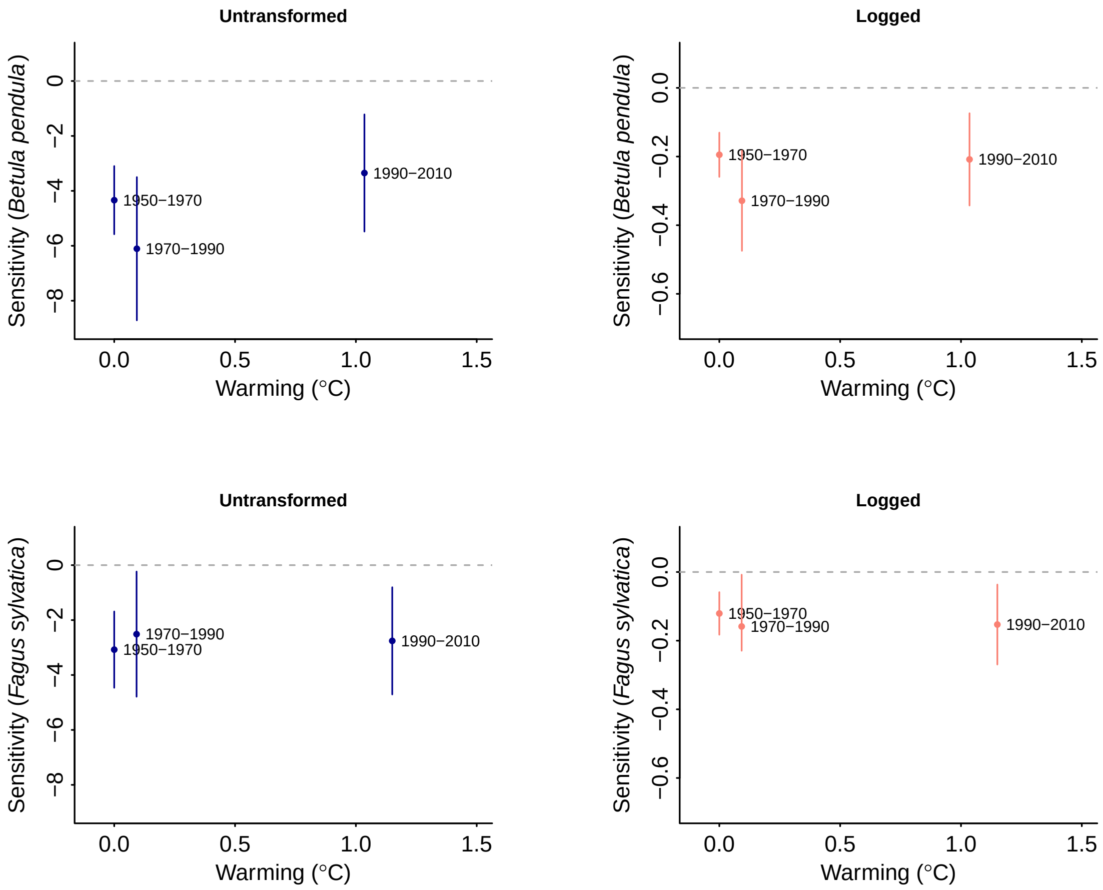


Figure S7: **Sensitivities from PEP725 data using 20 year windows of data** for two species (top – *Betula pendula*, bottom – *Fagus sylvatica*; all lines show 78% confidence intervals from linear regressions). Amounts of warming are calculated relative to 1950-1970 and we used only sites with leafout data in all years shown here. See Table S2 for further details.
